# Quantifying formations of Neutravidin clusters on biotinylated mixed silane Self-Assembled Monolayers

**DOI:** 10.1101/2022.03.21.485091

**Authors:** Ushnik Ghosh

## Abstract

Towards the goal of developing bio-chip / lab-on-a-chip substrates capable of performing highly specific bio-chemical reactions, Neutravidin binding to mixed Biotinylated Silane Self-Assembled Monolayers were studied using Confocal Fluorescence Light Microscopy. Non-specific bindings, specifically the formations of Neutravidin clusters, were quantified. Several experiments were conducted to determine the concentrations of Neutravidin necessary to not saturate surface binding to Biotinylated Self-Assembled Monolayers, determine the effectiveness of using FBS blocking buffers to reduce non-specific binding, optimize the repeatability of Neutravidin binding to Biotinlyated mixed Self-Assembled Monolayers with Silane-PEG-Biotin compositions ranging from 0 to 15%, and quantify background Neutravidin bindings and the corresponding formations of Neutravidin clusters to Self-Assembled Monolayers as Silane-PEG-Biotin percent compositions increase from 0 to 15%. The Neutravidin, Silane-PEG-Biotin, and Silane mPEG concentrations and ratios needed to develop homogeneous Neutravidin films, without the formations of clusters, on the Self-Assembled Monolayers have been determined.

## Introduction

Engineered Self-Assembled Monolayers composed of Alkanethiols on gold, or Silane functionalized PolyEthylene Glycol chains on glass, can be used to host embedded, nanostructured, supra-molecular assemblies such as proteins, glycoproteins, and polysaccharides on the monolayer surface as receptors to model and study ligand – receptor interactions [1]. Prior studies have demonstrated that it is possible to develop mixed Self-Assembled Monolayers, and modulate the composition and densities of the terminal ends of the alkanethiol or Silane-PEG molecules to design ligand – receptor chemistries [2,3]. The terminal ends of the alkanethiols or Silane-PEG chains can be chemically modified to host a wide array of receptors. The density of the receptors on the monolayer surface can be then be modulated by changing the ratios of the receptor terminated alkanethiols/Silane-PEG chains and non receptor terminated alkanethiol/Silane-PEG chains during the formation of the mixed Self-Assembled Monolayers.

Streptavidin binding properties to biotin terminated Self-Assembled Monolayers have been studied extensively in the literature. Biotin terminated mixed Self-Assembled Monolayers have been used with Streptavidin to host molecules such as DNA [4-7], thrombin [7], RGD [8], and multi-layer formations of bio-engineered flagella as protein nanotubes [9] for ligand - receptor interactions and kinetics studies. Streptavidin binding to Self-Assembled Monolayers with varying percentages of biotin have been quantified using Surface Plasmon Resonance spectroscopy (SPR) [4, 10-12], Quartz Crystal Microbalance with dissipation (QCM-D) [6, 12-15, 25], Ellipsometry [14, 15], Atomic Force Microscopy (AFM) [16, 17], and Fluorescence microscopy [18, 19].

The advantages of using Biotin – Streptavidin chemistry for functionalizing engineered Self-Assembled Monolayers are 1) Biotin and Streptavidin bind with a very high affinity and form one of the strongest non-covalent interactions known in nature, and 2) Streptavidin’s ability to bind with upto 4 biotin molecules, enabling the use of this molecule to host multiple bindings / reactions for complex chemical reactions. When used with Self-Assembled Monolayers, one binding site is occupied by the biotin terminated alkanethiol / Silane PEG chain, leaving 3 binding sites available for binding with 3 more biotinylated molecules. Thus the Biotin - Streptavidin complex has high versatility in that it can be used to host up to 3 different reactions to functionalize Self-Assembled Monolayers using the same chemistry.

For the development of a substrate that is capable of hosting ligand – receptor reactions with specificity, there are many properties of the Self-Assembled Monolayer that must be characterized. Previous studies have highlighted the importance of taking reaction environmental factors into account when designing ligand – receptor reactions, and have addressed factors such as steric hindrance to play a large role in the effectiveness of the interaction specificity. The use of homogeneous biotin-terminated Self-Assembled Monolayers are known to introduce steric hindrance between immobilized Streptavidin ligands during the binding process on the monolayer surface [21, 22]. The use of mixed Self-Assembled Monolayers, i,e. heterogeneous biotin functionalization, has been shown to reduce non-specific binding and steric hindrance of the Streptavidin molecules, and increase the quality of the Streptavidin film formation to the Self-Assembled Monolayer.

Poly-Ethylene Glycol (PEG) chains are widely incorporated into Self-Assembled Monolayer designs to reduce or eliminate non-specific adsorption of proteins [1 - verify, 23- 25]. The ability of the substrate to resist non-specific protein adsorption increases with the density of the Poly-Ethylene Glycol chains in the Self-Assembled Monolayer. Ellipsometry and SPR spectroscopy studies have demonstrated that the Poly-Ethylene Glycol chains in Alkane or Silane chains of Self-Assembled Monolayers play a key role in resisting non-specific adsorption of proteins [26]. Other factors such as solvent, concentration of Alkane or Silane chains, time length of immersion, and temperature also play a role in how well the Self-Assembled Monolayer can resist protein adsorption [1]. BSA, FBS, milk, and Tween-20 are commonly used blocking agents that can also be used in reactions to reduce or eliminate non-specific protein adsorption [27 -29]. The length of Alkane or Silane chains also play a large role in the prevention of non-specific protein adsorption onto the surface of Self-Assembled Monolayers. As the length of the Alkane / Silane chains increase, the Self-Assembled Monolayers assemble in a crystalline-like state and the lattice becomes less flexible, whereas Self Assembled Monolayers of shorter length assemble in a more disordered state. Highly structured Self-Assembled Monolayers are less likely to bend under their own weight, and subsequently are more resistant to non-specific protein adsorption to the surface [30-32].

Cluster formations of Streptavidin is a type of non-specific binding that has been observed in prior studies that have quantified the degree of non-specificity of Streptavidin binding to biotinylated Self-Assembled Monolayers [17, 33-34]. Azzaroni *et. al*. 2007 have quantified the formation of Streptavidin clusters on biotinylated Self-Assembled Monolayers in real-time using QCM-D [33]. Streptavidin molecules form a film on the biotin terminated Self-Assembled Monolayer after they are immobilized to the monolayer surface. Following maximum surface binding, the Stretavidin film re-organizes to become more stiff. Azzaroni *et. al*. 2007 have attributed the kinetics of the Streptavidin film reorganization in terms of the Lifshitz-Slyozov Law. Mir *et. al*. 2008 have used SPR spectroscopy to quantify the percentage of non-specific Streptavidin binding to biotinylated Self-Assembled Monolayers. In this experiment they have determined that 40% of the Streptavidin molecules have binding non-specifically to the biotinylated Self-Assembled Monolayers [34]. Lehnert *et. al*. 2011. has used AFM to image Streptavidin clusters on biotinylated silane-Self-Assembled Monolayers [17].

The formation of Streptavidin clusters can introduce challenges towards the development of substrates capable of hosting complex bio-chemical reactions with high specificity. Cluster formations of Streptavidin can increase the probability of non-specific bio-chemical interactions, and reduce the ability to host reactions with accuracy and precision. Neutravidin can be used as an alternative to Streptavidin towards efforts to reduce non-specific binding [35]. Neutravidin is a deglycosylated native avidin from egg whites, has excess carbohydrate removed that yields in a protein with a more neutral isoelectric point, and promotes to less non-specific binding. Removing the glycosylation of Streptavidin reduces carbohydrate-based lectin binding to undetectable levels without altering the biotin-binding affinity. Billah *et. al*. 2008 [36] has imaged Neutravidin binding to mixed biotinylated Self-Assembled Monolayers using Atomic Force Microscopy (AFM), and have observed Neutravidin cluster formations similar to Streptavidin cluster formations observed by Lehnert *et. al*. 2011 [17].

The literature cited have motivated this investigation to quantify Neutravidin cluster formations on biotinylated mixed Self-Assembled Monolayers. The experiments conducted in this study have characterized binding properties of Neutravidin to biotinylated Silane Self-Assembled Monolayers towards the goal of developing Neutravidin – Self-Assembled Monolayer substrates with minimum non-specific binding (Figure 1). Although the non-specific binding (cluster formations) between the Neutravidin molecules and biotinylated Self-Assembled Monolayers were still observed, this investigation has quantified increases of Neutravidin cluster formations that correspond with increases of background Neutravidin binding as the Silane-PEG-Biotin percentage compositions in the Self-Assembled Monolayers were increased. The Neutravidin molecules used in this study were conjugated with Oregon-Green-488 fluorophores, which enabled the investigator to quantify background Neutravidin binding and Neutravidin cluster formation using a Confocal Fluorescence Microscope.

**Figure 1:**
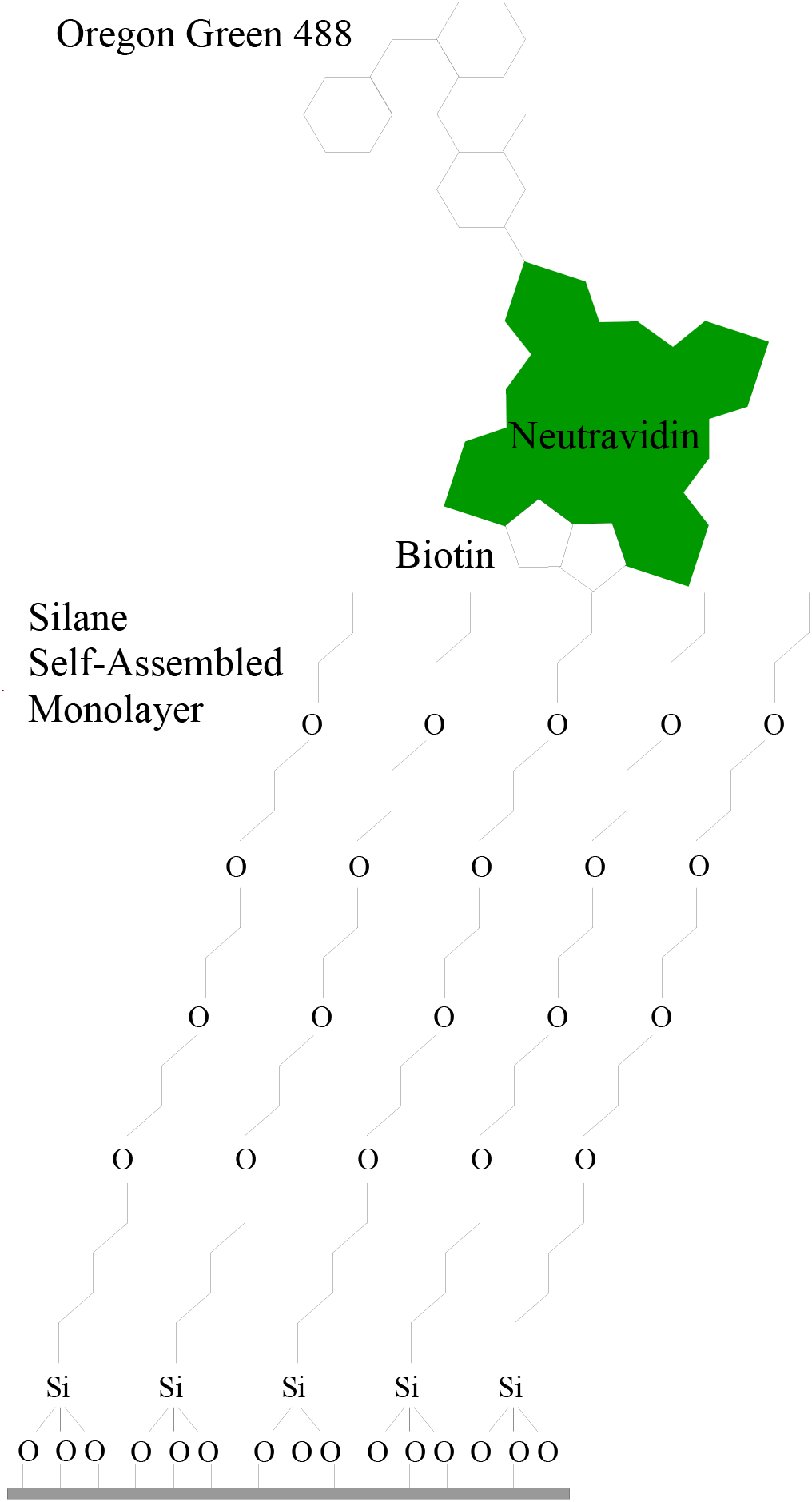
Mixed Self-Assembled Monolayer composed of Silane-mPEG and Silane-PEG-Biotin. Neutravidin molecules conjugated with Oregon-Green 488 fluorophores will be used to study Neutravidin binding to the biotinylated Self-Assembled Monolayers using a Confocal Fluorescence Light Microscope.

Previous studies have demonstrated that Streptavidin bindings to the surface of Self-Assembled Monolayers increase as surface biotinylation percentages increase from 0 to 20 percent, and the bindings decrease for biotinylation percentages from 20 to 100 percent [8, 10-11]. The background Neutravidin bindings and cluster formations to Silane Self-Assembled Monolayers were quantified at Silane-PEG-Biotin composition ranges of 0 to 15 percent, where the Neutravidin bindings are most responsive to biotinylation percentage compositions in the Self-Assembled Monolayers. The degree of Neutravidin clustering was quantified using the threshold function in the Image-J image processing software. Studying the Neutravidin bindings at this range enables the quantification of Neutravidin cluster formations that correspond with increases of background Neutravidin bindings as a function of surface biotinylation. The results from this study can be significant for future development efforts for homogeneous Neutravidin - biotinylated mixed Silane Self-Assembled Monolayer surfaces capable of hosting highly specific bio-chemical interactions.

This investigation has characterized the density, specificity, and homogeneity of biotin – Neutravidin binding to mixed biotinylated Self-Assembled Monolayers that can be used to host ligand – immobilized protein reactions. The density of Neutravidin binding to the Self-Assembled Monolayers were controlled by modulating the percent biotin composition in the mixed Self-Assembled Monolayers. However, gaining the ability to control the density of Neutravidin binding resulted in increases in Neutravidin cluster formations that introduced the challenge for developing homogeneous Neutravidin – Self-Assembled Monolayer substrates. The methods presented in this investigation have been optimized for the development of homogeneous Neutravidin – biotinylated Self-Assembled Monolayer substrates. The extent to which the density of Neutravidin binding to the mixed biotinylated Self-Assembled Monolayers can be modulated, and at what conditions the chemistry can be optimized to yield homogeneous substrates are discussed.

## Methods

### Materials

Glass bottom dishes were purchased from Mat-tek. Silane mPEG silane (5k) and Silane PEG Biotin (5k) were purchased from Nanocs. Neutravidin-Oregon green 488 fluorophores in PBS buffer (pH 7.4) was purchased from Thermo Fisher Scientific. PBS buffer (pH 7.4) was purchased from Thermo Fisher Scientific.

### Developing Self-Assembled Monolayers

Sterile glass bottom dishes were used to develop Silane Self-Assembled Monolayers. Silane mPEG silane (5k) and Silane-PEG-Biotin (5k) molecules were immersed in 95% Ethanol – 5% water solution at 13mg/ml concentration to form PEGylation solutions. The PEGylation solutions were put onto the glass bottom dishes, where Silane Self-Assembled Monolayers were formed in agitated conditions for 30 minutes in room temperature. The PEGylation solutions were then gently washed out, and the Self-Assembled Monolayers on the glass bottom dishes were rinsed 3 times with PBS buffer (pH 7.4). Throughout this paper the percent Silane-PEG-Biotin compositions are determined by the molecular weight ratio of Silane-PEG-Biotin used divided the total molecular weight of Silane-PEG-Biotin and Silane-mPEG molecules used to create the Self-Assembled Monolayer.

### Binding Neutravidin to Silane-PEG Self-Assembled Monolayers

Neutravidin-Oregon Green 488 molecules were pipetted onto the center of the Self-Assembled Monolayers at 4, 8, 16, 32, 128, 256, 1024, and 2048 µg/ml concentrations and were allowed to bind to the Silane-PEG-Biotin molecules for 30 minutes. Neutravidin-Oregon Green 488 solution concentrations changed with PBS buffer dilutions. 5% FBS, 5% BSA, or 5% milk blocking buffers were added to the Neutravidin-Oregon Green 488 in PBS solution for experiments using additional measures to reduce non-specific binding. The glass-bottom dishes with Neutravidin bound Self-Assembled Monolayers were then gently rinsed 3 times with PBS buffer (pH 7.4) to wash off excess Neutravidin. Glass-bottom dishes with Self-Assembled Monolayers were then flooded with PBS buffer to prevent protein denaturation during the imaging process.

Confocal Fluorescence Imaging: Glass-bottom dish samples were imaged using an Olympus FV1000 Confocal Microscope.. Neutravidin-Oregon Green 488 molecules were imaged using 473 nm excitation light. Samples were imaged at three distinct locations throughout the Neutravidin film on the Self-Assembled Monolayer using a 10X objective lens.

Fluorescence Intensity Analysis: Confocal microscopy images of Neutravidin-Oregon Green 488 bound Self-Assembled Monolayers were analyzed using Image-J Fiji. Background fluorescence intensities were determined by measuring the average brightness of the image pixels throughout the images. Neutravidin cluster fluorescence intensities were quantified using the threshold function to measure the number of pixels that were in the top 4.9% of the intensity distribution histogram. This threshold was applied uniformly for all images.

## Results

Four experiments were conducted to determine the conditions necessary to reduce or eliminate Neutravidin cluster formations towards the efforts of developing homogeneous and uniform Neutravidin binding to Self-Assembled Monolayers. The minimum Neutravidin concentrations necessary to observe specific binding yields were determined in the first experiment. Non-specific binding was minimized by determining the the minimum concentration of Neutravidin that was necessary to observe specific binding without the use of blocking buffers. The use of blocking buffers to effectively remove or reduce non-specific Neutravidin binding to Self-Assembled Monolayers are demonstrated in the second experiment. The Neutravidin concentrations and blocking concentrations necessary to yield minimum non-specific binding were determined in the first two experiments. Neutravidin bindings to mixed Biotinylated Silane Self-Assembled Monolayers were quantified as Silane-PEG-Biotin compositions were increased from 0 – 15% in the third experiment. The fluorescence intensities of blank substrates with and without immobilized Neutravidin were also quantified as positive and negative controls. Background Neutravidin binding and the corresponding Neutravidin cluster formations were imaged and quantified as Silane-PEG-Biotin compositions on the mixed Biotinylated Silane Self-Assembled Monolayers were increased from 0 to 15% in the fourth experiment.

### Determining the concentration of Neutravidin molecules that does not saturate the binding to Self-Assembled Monolayers

A thorough experiment is conducted where Self-Assembled Monolayers with varying Silane-PEG-Biotin and Silane mPEG ratios were developed to determine the concentration of Neutravidin that is necessary to see a density response in Neutravidin and Silane-PEG-Biotin binding (Figure 2). The goal of this experiment was to determine the saturation point of Neutravidin binding to the Self-Assembled Monolayers. Saturating the surface of the Self-Assembled Monolayers with Neutravidin may cause a Neutravidin monolayer formation that does not reflect biotin – Neutravidin specific binding. Determination of the Neutravidin concentrations that no longer saturate the Self-Assembled Monolayers is a necessary step to identify the conditions that will yield highly specific Biotin – Neutravidin bindings. Neutravidin molecules were pipetted at 4, 8, 16, 32, 128, 256, and 1024 µg/ml concentrations onto Self-Assembled Monolayers with 0%/100%, 25%/75%, 50%/50%, 75%/25%, and 100%/0% Silane-PEG-Biotin/Silane mPEG ratios. The average Neutravidin bindings to Self-Assembled Monolayers decrease as the Neutravidin concentrations are reduced from 2048 to 16 µg/ml (Figure 2). A one-way ANOVA test has determined that the concentration of immobilized Neutravidin molecules yield in statistically significant differences for quantified fluorescence intensities, with a p-value of 0.0002. The average bindings decrease significantly when Neutravidin molecules are immobilized at 4 and 8 µg/ml concentrations. This is an indication that Neutravidin bindings at 4 and 8 µg/ml concentrations do not saturate the surface of the Self-Assembled Monolayers and are likely to be more specific. Neutravidin molecules are immobilized at concentrations of 8 µg/ml for the remainder of the experiments conducted in this investigation. It is also interesting to note that there is no observable trend as percent Silane-PEG-Biotin compositions are modulated for each concentration of Neutravidin immobilizations. This reflects the need for additional means to control non-specific binding.

**Figure 2:**
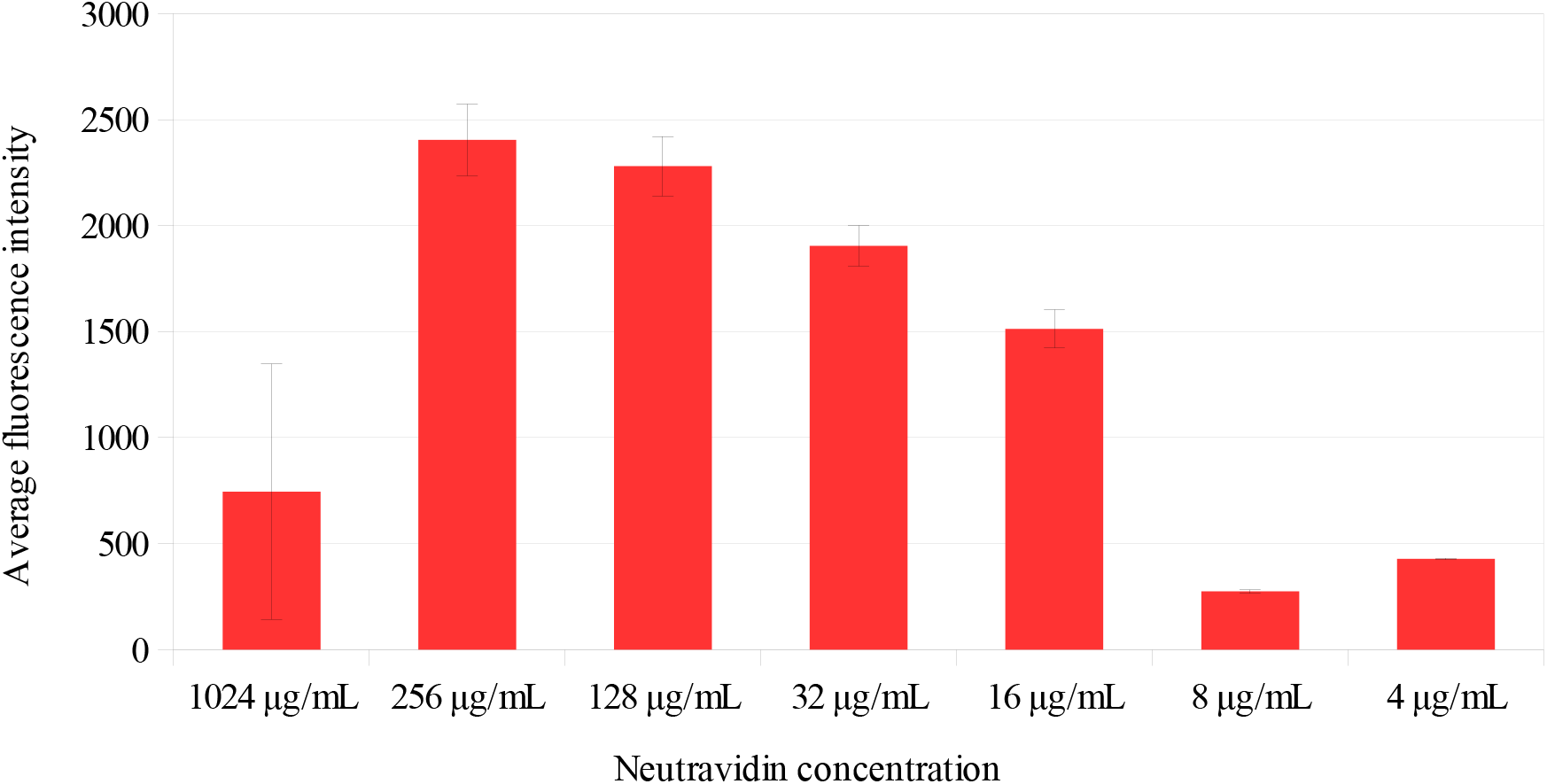
Determining the concentration of Neutravidin molecules that does not saturate the binding to Self-Assembled Monolayers. Fluorescence intensity of varying Neutravidin-Oregon Green 488 concentrations to 0-100% [25%] biotin Self-Assembled Monolayers. Fluorescence intensity does not change significantly when Neutravidin concentrations of 1024-16 µg/ml are pipetted onto the Self-Assembled Monolayer. Neutravidin at concentrations of 8-4 µg/ml, experience significant decrease in average binding. Error bars represent the standard deviation of the samples.

### Blocking non-specific Neutravidin binding to Silane Self-Assembled Monolayers

In addition to using PEG modified Silane chains to construct the Self-Assembled Monolayer, the non-specific Neutravidin adsorption and formation of clusters to the substrates were reduced with the use of FBS, BSA, or Milk blocking buffers. FBS, BSA, and Milk blocking buffers were used to successfully eliminate non-specific binding in 100% Silane mPEG Self-Assembled Monolayers and in negative control glass substrates with no Self-Assembled Monolayers (Figure 3). The results from this experiment and the previous experiment were used as preparatory measures for minimizing non-specific Neutravidin binding to the Self-Assembled Monolayers, and will enable the study and quantification of Neutravidin cluster formations with greater control. It is important to note that Neutravidin cluster formations can be observed on the 100% Silane mPEG Self-Assembled Monolayers samples and not on the glass substrate controls. This is an indicate that Neutravidin cluster formation is a property of Neutravidin – Self-Assembled Monolayer interactions.

**Figure 3:**
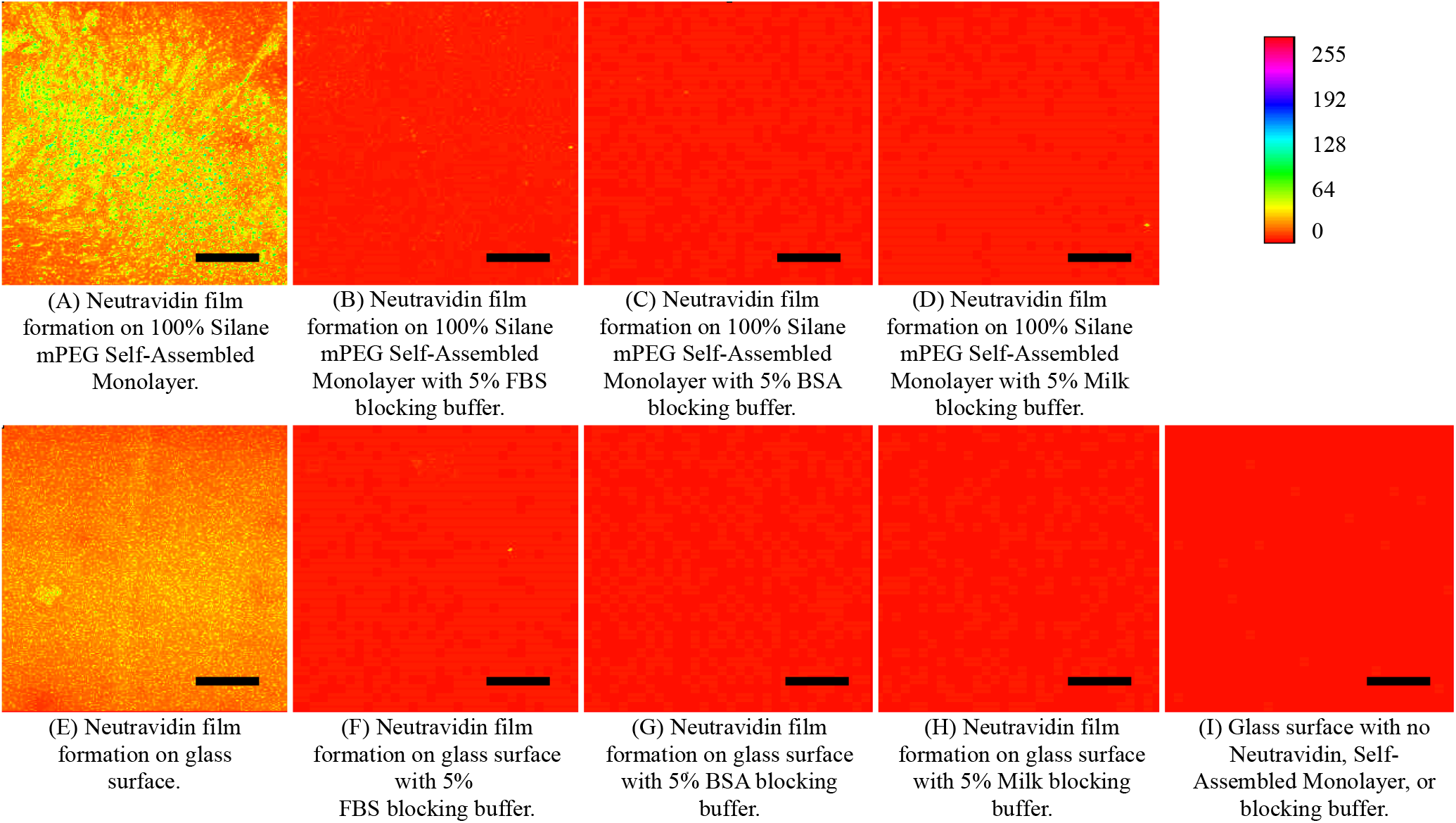
Figure 3.A shows the formation of Neutravidin clusters on a 100%mPEG-Silane Self-Assembled Monolayer. Figures B. - D. demonstrate that non-specific Neutravidin cluster formation on the 100% mPEG-Silane Self-Assembled Monolayer can be prevented by using 5% FBS (Figure B), 5% BSA (Figure C), or 5% Milk (Figure D) blocking buffers. Figure E. shows Neutravidin binding to glass substrate Figures F - H demonstrate that FBS, BSA, or Milk blocking buffers also prevent non-specific Neutravidin cluster formation on glass substrates. Figure I. Shows the fluorescence of a glass substrate without a Self-Assembled Monolayer or Neutravidin. Scale bars represent 100µm.

### Determining Variability of Neutravidin fluorescence intensity by modulating Silane-PEG-Biotin percentage composition in mixed Self-Assembled Monolayer

An experiment was conducted to identify the ideal molecular Silane-mPEG and Silane-PEG-Biotin concentration ratios needed to develop reapeatable, homogeneous Neutravidin monolayers. The graphs from Figure 4 show three experiments where the bindings of immobilized Neutravidin molecules with 5% FBS blocking buffer were quantified at 8µg/ml concentrations to Self-Assembled Monolayers with 0%, 5%, 10%, and 15% Silane-PEG-Biotin compositions. The data points of this graph were collected by quantifying the Neutravidin - Oregon-Green 488 bindings to 36 Self-Assembled Monolayers over three experiments, with measurements from three distinct locations from each Self-Assembled Monolayer. A one-way ANOVA test was used to test if the percent Silane-PEG-Biotin composition has a statistically significant Neutravidin binding response, and has yielded a p value of 0. The x-axis label denotes the percentage of Silane-PEG-Biotin composition in the respective sample, and the y-axis denotes the quantified Neutravidin fluorescence intensity value calculated using the intensity analysis tool of the Image-J image processing software. The experiments have similar trends for Neutravidin binding yields. As expected, Neutravidin bindings increase as surface Silane-PEG-Biotin percentage composition in the Self-Assembled Monolayers increase for all three experiments. The fluorescence of blank samples (0% Silane-PEG-Biotin, 0% Silane-mPEG) with and without immobilized Neutravidin has also been quantified, and very little difference in the fluorescence intensity is observed. These negative controls were used to verify that the Neutravidin does not bind non-specifically to glass-substrates that does not have any Silane-PEG-Biotin, and very that there is little background fluorescence.

**Figure 4:**
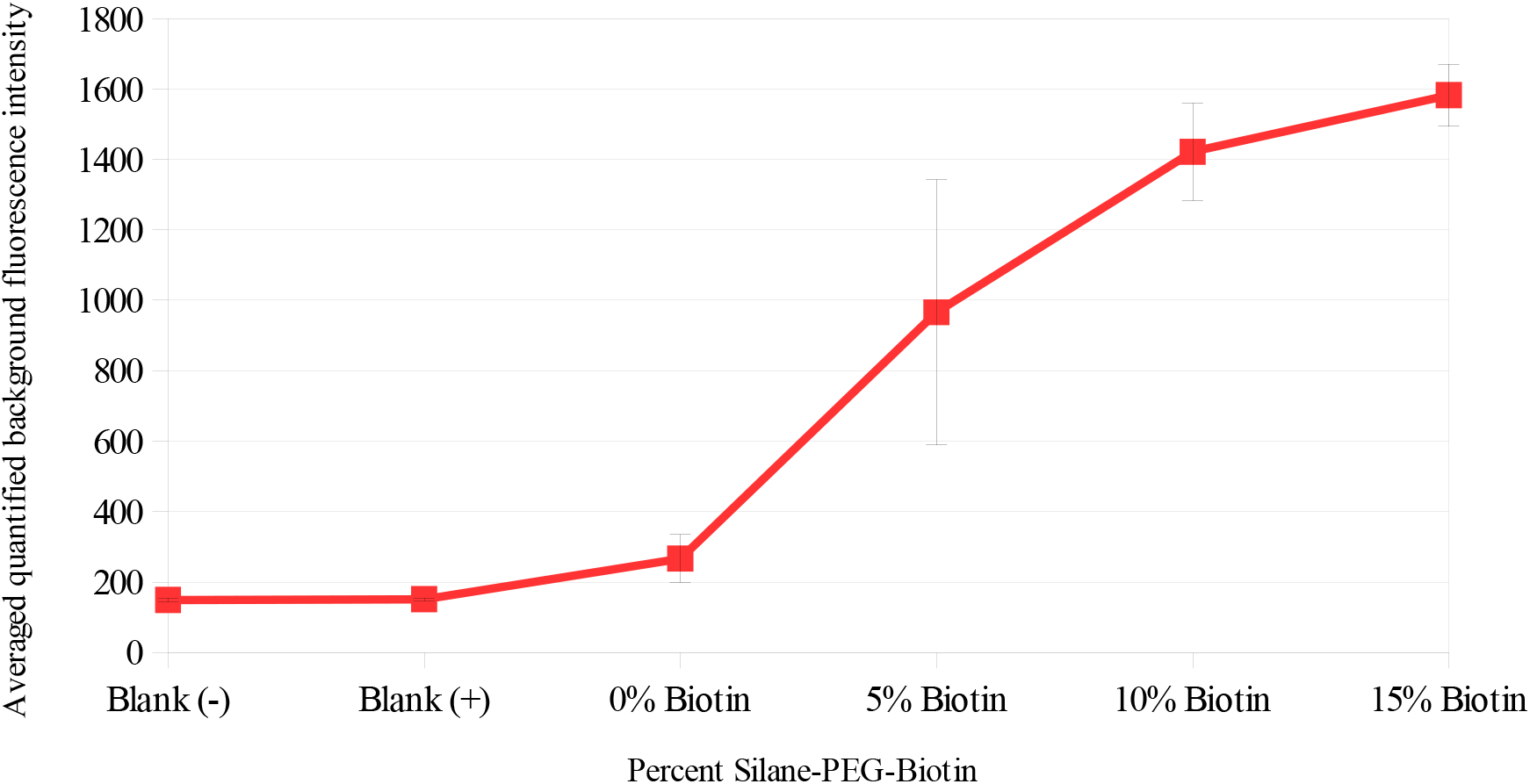
Fluorescence intensities of Neutravidin-Oregon Green 488 (immobilized at 8µg/ml concentrations) binding to 0%, 5%, 10%, 15% Silane-PEG-Biotin Self-Assembled Monolayers across 3 experiments. Fluorescence intensities of control samples without Neutravidin or Self-Assembled Monolayers is shown as Blank(-). Fluorescence intensities of Neutravidin binding to blank samples without Self-Assembled Monolayers is shown as Blank(+).

### Background Neutravidin bindings and corresponding cluster formations to biotinylated Self-Assembled Monolayers

The goal of the fourth experiment was to determine the conditions that are needed to develop a Neutravidin film with minimum cluster formation. Using the results from the previous sections, a set of Self-Assembled Monolayers were developed to study Neutravidin bindings and the corresponding cluster formations. Figures 5.B – 5.E show Neutravidin background fluorescence and the corresponding increase in cluster formation as percent Silane-PEG-Biotin within the Self-Assembled Monolayers increase. Neutravidin-Oregon Green 488 molecules at 8µg/ml concentrations with 5% FBS blocking buffer were pipetted onto and rinsed off of the Self-Assembled Monolayers with Silane-PEG-Biotin percent compositions ranging from 0% to 15%. Blank glass substrates with no Self-Assembled Monolayers, that have had Neutravidin pipetted onto the surface and then rinsed off with PBS, do not have any background Neutravidin bindings or cluster formations (Figure 5.A), confirming that the FBS buffer successfully blocks non-specific binding. 100% mPEG-Silane (0% Biotin) Self-Assembled Monolayers with blocking buffer yield a very low density, homogeneous Neutravidin film (Figure 5.B.), indicating that there is a very small amount of Neutravidin non-specifically binding to the mPEG-Silane molecules. As the percentage of Silane-PEG-Biotin composition in the Self-Assembled Monolayers increase, the background fluorescence, and the corresponding cluster formations increase (Figure 5.C.-5.E.). Neutravidin binding to Self-Assembled Monolayers with 5% Silane-PEG-Biotin composition yield in uniform and homogeneous Neutravidin films, with little cluster formation. For Self-Assembled Monolayers with higher percentages of Silane-PEG-Biotin compositions, there is a noticable increase of Neutravidin cluster formations. Figure 6 shows the quantified fluorescence intensities of background Neutravidin binding, as well as the quantified fluorescence intensities of the corresponding Neutravidin cluster formations, for the same experiment. Each bar on the graph is the average of three quantified Neutravidin fluorescence intensity values from three distinct regions on a Self-Assembled Monolayer. The quantified fluorescence intensity analysis shows an increase in Neutravidin cluster formations as background Neutravidin bindings increase. Paired-t-tests were used to determine whether the difference between background and cluster fluorescence intensity values are statistically significant. The p-values were 0.0082, 0.2038, 0.0076, and 0.0007 for Self-Assembled Monolayers with 0%, 5%, 10%, and 15% Silane-PEG-Biotin compositions.

**Figure 5:**
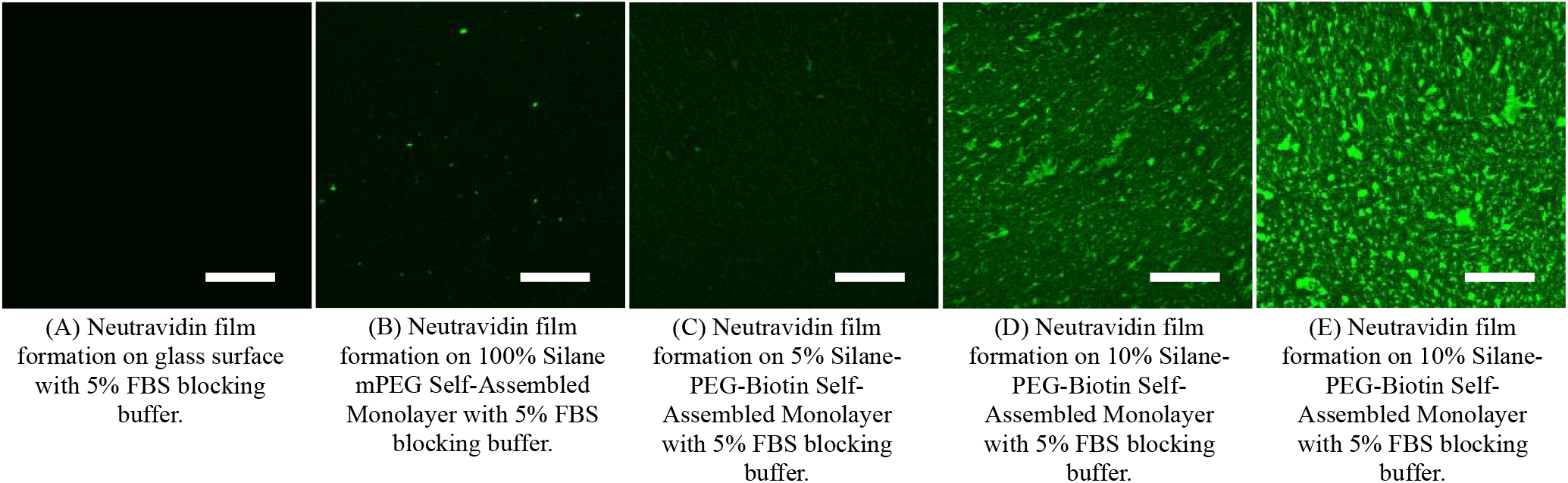
Figures A to E shows Neutravidin-Oregon Green 488 (8µg/ml) binding to the surface of Self-Assembled Monolayers as the percentage of biotin increases from 0%-15%. Figure A Oregon-Green 488 fluorescence of a glass substrate that has been bound with Neutravidin. Figure B shows the background fluorescence of a 100% Silane mPEG (0% Biotin) Self-Assembled Monolayer. Figure C shows the background fluorescence of a 5% Silane-PEG-Biotin Self-Assembled Monolayer. Figure D shows the background fluorescence of a 10% Silane-PEG-Biotin Self-Assembled Monolayer. Figure E shows the background fluorescence of a 15% Silane-PEG-Biotin Self-Assembled Monolayer. Scale bars represent 100µm.

**Figure 6:**
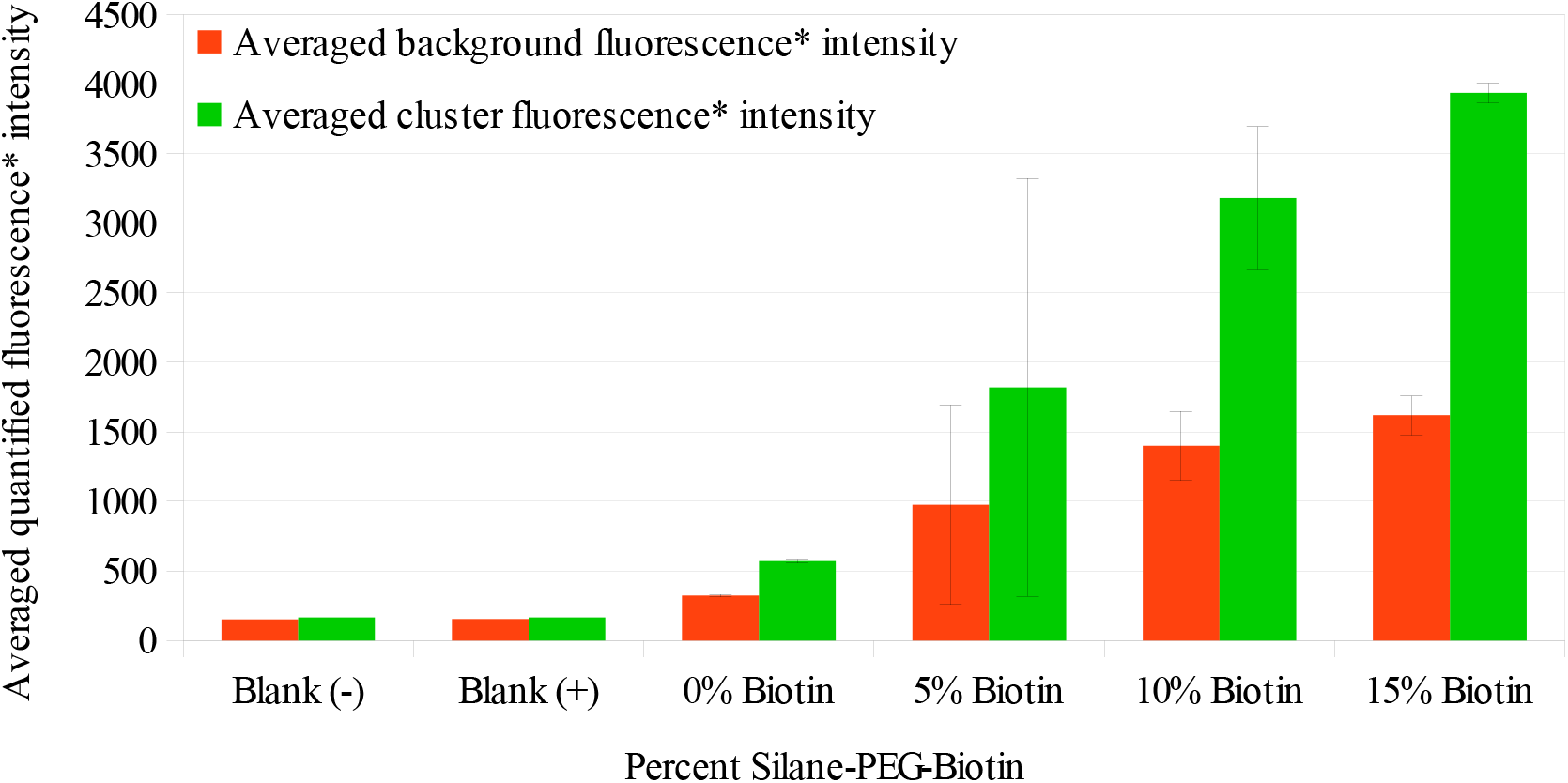
Quantified average background Neutravidin Oregon-Green 488 fluorescence intensities (blue) and corresponding quantified average Neutravidin cluster fluorescence intensities (red) for Self-Assembled Monolayers with 0%, 5%, 10%, and 15% Silane-PEG-Biotin compositions. The fluorescence intensities of background Neutravidin and clustered Neutravidin on blank substrates (with no Self-Assembled Monolayers are also shown). Each bar on the graph is the quantified Neutravidin fluorescence intensity values that have been averaged from 3 distinct regions on a Self-Assembled Monolayer averaged for 3 Self-Assembled Monolayers. The error bars represent the standard deviation values for the averaged quantified fluorescence intensity values.

## Discussion

Self-Assembled Monolayers can be functionalized with Neutravidin to host a wide array of complex bio-chemical interactions towards the goals of studying ligand – receptor interactions. The design of Self-Assembled Monolayers and the surface functionalization play key roles in the effectiveness for modeling ligand – receptor interactions. It is necessary to characterize reactive functional-group concentrations and homogenuity during the design and development of bio-chip / lab-on-a-chip substrates for goals to maximize binding interaction specificity. Towards this goal, this investigation has studied the formations of Neutravidin clusters on Biotinylated Mixed Silane Self-Assembled Monolayers. The results from the final experiment have demonstrated formation of Neutravidin clusters is directly dependent on the background Neutravidin binding to the Biotinylated Self-Assembled Monolayers. Results from the second experiment show that the formation of Neutravidin clusters can also be observed in Self-Assembled Monolayer with 100% mPEG-Silane composition, but not on glass substrates with no Self-Assembled Monolayers. It is possible that the formation of Neutravidin clusters is a property of the Silane-PEG-Biotin compositions, as well as, the property of Silane-mPEG compositions of the Self-Assembled Monolayers. A good future study would be to discern whether the formation of Neutravidin clusters are a function of surface Biotin percentage composition or the property of interactions with other Self-Assembled Monolayer molecules such as Poly-Ethelene Glycol. Discerning the origins of the Neutravidin clusters can further efforts towards the design and development of Self-Assembled Monolayer based bio-chip / lab-on-a-chip substrates capable of modeling ligand – receptor interactions with high specificity.

## Notes

### Competing Interest Statement

The authors have declared no competing interest.

### Summary of Updates

The removed second author's name was flickering even though it was painted over. I have editted the name out with adobe pro, and I think that will fix this problem.

